# Assessing the impact of pedigree quality on the validity of quantitative genetic parameter estimates

**DOI:** 10.1101/2022.11.03.514896

**Authors:** Walid Mawass, Emmanuel Milot

## Abstract

Investigating the evolutionary dynamics of complex traits in nature requires the accurate assessment of their genetic architecture. Using a quantitative genetic (QG) modeling approach (e.g., animal model), relatedness information from a pedigree combined with phenotypic measurements can be used to infer the amount of additive genetic variance in traits. However, pedigree information from natural systems is not perfect and might contain errors or be of low quality. Published sensitivity analyses revealed a limited impact of expected error rates on parameter estimates. However, natural systems will differ in many respects (e.g., mating system, data availability, pedigree structure), thus it can be inappropriate to generalize outcomes from one system to another. French-Canadian (FC) genealogies are extensive and deep-rooted (up to 9 generations in this study) making them ideal to study how the quality and properties (e.g., errors, completeness) of pedigrees affect QG estimates. We conducted simulation analyses to infer the reliability of QG estimates using FC pedigrees and how it is impacted by genealogical errors and variation in pedigree structure. Broadly, results show that pedigree size and depth are important determinants of precision but not of accuracy. While the mean genealogical entropy (based on missing links) seems to be a good indicator of accuracy. Including a shared familial component into the simulations led to on average a 46% overestimation of the additive genetic variance. This has crucial implications for evolutionary studies aiming to estimate QG parameters given that many traits of interest, such as life history, exhibit important non-genetic sources of variation.

## Introduction

Advancing the predictive study of evolutionary dynamics in the wild rests on the proper estimation of genetic parameters underlying phenotypic variation, such as the additive genetic (co)variance. Over the last quarter of a century, the field of quantitative genetics (henceforth QG), drawing from the early work of animal breeders (Lush 1937; Henderson 1975) and pioneer work extending it to natural populations (Shaw 1987), popularized the application of pedigree-based methods to estimate genetic parameters in the wild (e.g., Lynch & Walsh 1998; Kruuk et al. 2000; Charmantier et al. 2014). These methods can exploit all known filial relationships between individuals in the pedigree to dissect phenotypic variation into its genetic and other components (e.g., environmental, maternal; Kruuk 2004; Wilson et al. 2010; Walsh & Lynch 2018). One popular method is to fit to the data the ‘animal model’, a special case of generalized linear mixed-effect models that can deal with unbalanced pedigrees typically associated with natural populations. Various implementations of the animal model are now available (Gilmour et al. 2002; Hadfield 2010; De Villemereuil et al. 2013, 2016; Rousset 2017; see van Benthem 2017 for a review) and have been used to study microevolutionary processes in nature (e.g., Foerster et al. 2007; Wilson et al. 2006, 2007; Milot et al. 2011; Fisher et al. 2019).

Pedigrees provide information on the expected pairwise relatedness between individuals, as measured by the kinship coefficient *Θ*_*ij*_, namely the probability that a gene copy randomly drawn from individual *i* is identical-by-descent to a copy randomly drawn from individual *j* for the same gene. This information is used in conjunction with phenotypic measurements to estimate QG parameters, based on the principle that the degree of additive genetic determination of a trait determines the phenotypic resemblance between relatives. How good is parameter estimation strongly depends on the information contained in the available pedigree, which can vary much among natural, hence uncontrolled, populations (Bonnet et al. 2022). The validity of a method is its capacity to correctly measure what it should measure. In the context herein, we define it as the capacity of a QG method to estimate parameters of interest (i.e., genetic) with acceptable accuracy and precision, and over the range of their potential values, given the pedigree. In other words, is it valid to fit animal models to this particular pedigree? This operational definition is tailored to our goal, which is to study how good pedigrees (with given attributes) can be for the estimation of genetic parameters (as opposed to how good the animal model is in general, i.e., across pedigrees). While what is considered as “acceptable accuracy and precision” may vary among people or contexts, for the purpose of this study we will consider them as acceptable when they allow the reaching of correct qualitative conclusions.

Pedigree attributes influencing the validity of a QG analysis can be roughly divided into two categories: those related to data, such as completeness and errors, which reflect our state of knowledge about the true pedigree; those related to topology, which result from the biology of the population itself (see Table 1 for a summary of concepts). Regarding the first category, one assumption underlying QG is that, unless specified explicitly in the model structure, a pedigree provides errorless values of pairwise relationships between individuals. For it to be the case requires three conditions: i) genealogical links between parents and offspring are correct; ii) no links are missing between individuals who appear in the pedigree, such that all true genealogical paths connecting a pair of phenotyped individuals are known (relative to the basal population); iii) pedigree-based relatedness matches genetic relatedness, recalling that a pedigree provides expected (mean) relatedness values, not accounting for the variance introduced by Mendelian segregation and recombination. Thus, obtaining errorless relatedness values is unrealistic (Hill & Weir 2011), both for captive stocks in animal breeding programmes (Visscher et al. 2002) and natural populations (Speed and Balding 2015). Pedigree errors and incompleteness generally reduce the extent to which relatives appear to resemble each other, causing genetic parameter underestimation. For example, in captive breeds of Hereford cattle, estimates of heritability in two traits, birth weight and weaning weight, decreased linearly as the rate of sire misidentification increased (Senneke et al. 2004).

**Table 1.** Table showing the key factors which affect pedigree quality and performance.

| Attribute of | Attribute name | Description | Notes |
| --- | --- | --- | --- |
| True pedigree | Depth | Number of generations separating the current (most recent) cohort and population founders. | From a QG perspective, founders are those who represent the basal population. |
|  | Pedigree structure | Differing mating systems and reproductive strategies lead to contrasting pedigree structures, a large pedigree with a short generation time vs smaller pedigree with a longer generation time | Different pedigree structures perform better for specific estimations. For example, pedigrees from polygynous species are good for estimating shared environmental effects, while pedigrees from long-lived species perform well in studies of genotype-by-environment interactions |
| Data-reconstructed (or empirical) pedigree | Quality | In a broad sense, quality means how close is the reconstructed pedigree to the true, but unobserved, pedigree. | Quality is determined by attributes such as sample size, completeness, depth, and the occurrence of genealogical errors. |
|  | Sample size | Number of individuals who are known/used to build the pedigree. | Sometimes reported per generation. |
| | Completeness | Ratio of number of known ancestors to the number of expected ancestors of the sampled individuals. | A complete pedigree contains ancestors for a focal individual, where $g=0$ designates its own generation, $g=1$ that of its parents, and so forth. E.g., if $G=3$, then the individual has 14 ancestors<sup>a</sup>. A pedigree lacking the identity of 4 of them would have a completeness value of 0.71. |
| | Depth | Number of generations covered by the reconstructed pedigree (e.g., average number of generations separating individuals at lineage tips from their ancestors at lineage bases) | Balloux et al. (2004) consider that when depth is $< 5$ generations, and depending on mating system and population structure, a large proportion of variance in parameters such as coancestry, relatedness, and inbreeding coefficient, won't be captured. |
|  | Phenotypic coverage | The proportion of individuals in the pedigree for which phenotypic measurements are available. | Individuals with no measurements still contribute information to QG analyses when they lie on a genealogical path linking 2 individuals with measurements. However, genetic parameter accuracy and precision should increase with increasing phenotypic coverage, especially in cases where other sources of variation are present such as shared familial environments which require at least two or more sibs be phenotyped |
|  | Genealogical error | An incorrect link between two related individuals | This type of error occurs due to over-linkage between individuals (i.e., wrong biological parental assignments). It can result from hidden adoptions by social (non-biological) parents or false parentage declarations, for example. Genealogical errors can occur also from under-linkage which refers to missing parental assignments due to information that is lacking or unclear on historical acts |
|  | Pedigree accuracy | The accuracy of pedigree links which declines with the presence of errors caused by extra-pair paternities and parental misassignment | Link errors are expected to lead to downward-biased estimations of heritability (or additive genetic variance), i.e., lower precision. Inaccurate pedigree links can cause environmental covariance to be misinterpreted as genetic covariance. This is most likely in cases of small sample size and highly heritable traits. However, according to Charmantier & Réale (2005), animal model heritability estimates are robust when EPP rates are lower than 20% |
|  | Entropy | This factor also relates to data availability. It is the proportion of missing links or missing parentage assignment to the total pedigree links. It provides a measurement of how sparse the information is on the ancestral generations | Lack of information on parental assignment and pedigree links differ from inaccuracy in links in that it might not lead to downward-biased estimations and imprecise measurements if sample size is large enough, but it is expected to negatively impact the accuracy of estimates |

<sup>a</sup> When inbreeding occurs, these 14 ancestors need not be 14 different individuals. For example, one could be “twice” the great-grand-mother (i.e., through 2 genealogical paths) of an offspring born to a mating between cousins.

Pedigree topology reflects the properties of the natural population *per se*, such as its mating system, life history, and vital rates. For example, a polygynous population will exhibit a greater variance in male reproductive success than a monogamous population (all else being equal), which will lead to different kinship structure in their pedigree and, consequently, different amount of information available to infer parameters. In addition, it will not be possible to separate sire and dam effects in a monogamous species (unless monogamy is sequential, in which case information to measure these effects may still remain quite limited).

Sensitivity and power analysis are two means by which one can assess the adequacy of a pedigree for QG (as per Morrissey et al. 2007). Sensitivity consists in determining how parameter estimates change when genealogical errors are deliberately introduced in the pedigree. Power analysis measures the capacity to recover from QG analyses preset genetic (co)variance values used to simulate breeding values along the empirical pedigree. For example, Charmantier & Réale (2005) and Firth et al. (2015) conducted simulations on social pedigrees from blue tits (*Parus caeruleus*) and great tits (*P. major*; data from Firth et al. 2015) and showed that underestimation of heritability in body mass and tarsus length was fairly limited when extra-pair paternity (EPP) rates were up to 20% (note, however, that traits expressed in a single sex could react differently when genealogical errors involve mostly one parental sex, as in the case of EPPs). Bérénos et al. (2014) used three pedigrees with increasingly accurate estimates of relatedness (pedigree constructed from field observations; pedigree constructed from specific markers; pedigree constructed using whole-genome information) of an unmanaged Soay sheep population to show that field observation-based pedigree performed as well as SNP-based and whole-genome pedigree in terms of heritability estimates.

The impact of pedigree quality and topology (Table 1) on the validity of QG parameter estimates has received limited attention in evolutionary genetics. Only a minority of empirical studies that fitted animal models to wild population data appear to quantify how good their pedigrees are for QG analyses. In a quick informal survey of studies published over the past 10 years (*n*=30), we found only ∼⅓ of them performed a formal sensitivity analysis using simulations. Among the remaining ⅔, some studies limit their consideration of sensitivity issues to citing other studies, such as those aforementioned. This is of course of relevance, e.g. for result interpretation purposes, but does not amount to a quantitative assessment of the specific pedigree at hand. In addition, sensitivity analyses have mainly focused on genealogical link errors, such as those due to EPPs in social pedigrees, but less on missing links (i.e. pedigree completeness). Though both factors will generally decrease the apparent phenotypic resemblance between relatives, their quantitative impact on estimation can differ. Actually, Morrissey et al. (2007) showed that erroneous genealogical links cause a greater underestimation of QG parameters than missing links. Wolak & Reid (2017) found that parameters estimated with animal models will be biased when parentage links are missing in a systematic way with respect to genetic or phenotypic values of population founders. Moreover, pedigree errors or an incomplete pedigree can lead to a downward bias in the estimation of maternal effects, which are a relevant source of phenotypic similarity between relatives (Wilson et al. 2010), as well as of heritability, as aforementioned. All these issues relate to the problem of parameter identifiability in statistical models with complex variance structures (Bolker 2008). So far, it is still unclear how pedigree topology and attributes impact the identifiability of variance components in animal models (Bourret et al. 2017).

In this study we used socially-constructed human pedigrees based on the genealogies of French-Canadian (henceforth FC) historical populations provided by the BALSAC database (https://balsac.uqac.ca/), which span all Québec regions since their colonization by European settlers. FC populations offer two advantages to explore the relationship between pedigree attributes and genetic parameter estimation. The first one is the possibility to separate the huge Québec-wide pedigree into multiple sub-pedigrees with contrasting attributes, corresponding to several geographic subpopulations, and compare them in our analyses. The second advantage is that we have good estimates of genealogical error rates from previous gene-dropping simulations (Doyon et al. *in prep*.). Our objectives are: 1) to assess how pedigree attributes (i.e., sample size, depth, etc.) impact the validity of QG estimates; 2) evaluate the impact of different model parameterization and variance structures on the validity of QG estimates.

## Materials and Methods

### Pedigree quality

The sizable variation in the properties of different species and populations results in a diversity of genealogical topologies in nature, some of which being more tractable than others by researchers. Herein *pedigree quality* refers to how close is the data-reconstructed pedigree to the true (but unobserved) pedigree. It is determined by attributes such as sample size, depth, genealogical error rate, and completeness (Pemberton 2008; Table 1). A pedigree may be of low quality because it lacks depth relative to the basal population or contains a high number of parental assignment errors, for example. An incorrect link can contribute to parameter underestimation in two ways that mirror each other: i) it may wrongly connect an individual A to another individual B, while both are in reality little related to each other. As a consequence, for a given value of heritability, the pair will exhibit no or limited phenotypic resemblance due to IBD genes, relative to what would be expected from their presumed but wrong genealogical relationship; this problem also extends to pairwise comparisons between A and other individuals who are related to B; ii) it will cause the analysis to ignore a true link (and resemblance) between the same individual A and others with whom he/she is more related than apparent in the erroneous pedigree. In certain species, it can be difficult to establish offspring-father links due to extra-pair paternities (EPPs) or because of limited paternal care reducing the detectability of fathers. In other species, parentage is practically intractable in the field for both parents and it may be necessary to perform genetic assignment to parental pairs (see Perrier et al. 2013 for an example in a salmonid population).

Precision of QG estimates is impacted by pedigree depth, sample size, and the complexity of the genetic architecture underlying the focal trait (Morrissey et al. 2007; de Villemereuil et al. 2013). Between two pedigrees of identical size but differing by their degree of connectivity, the one with a weaker connectivity will provide less precise parameter estimates (Wilson et al. 2010). Accuracy is the proximity between the estimated value and the true value. All pedigrees, even those from controlled animal breeding designs, are likely to contain link errors affecting the accuracy of estimation (Charmantier & Réale 2005; Morrissey et al. 2007; Firth et al. 2015).

A pedigree said to be complete over *G* generations is one that contains all ancestors of the focal (e.g., phenotyped) individuals back to the first generation covered, as well as no missing genealogical links. For example, certain individuals have no known parentage because either i) they are founders or immigrants or ii) they were never successfully linked to possible parents in the reconstruction of the genealogy. Pedigree completeness varies when different lineages ascending from a focal individual are not uniformly covered (for example, when both paternal grand-parents are known for an individual but one only maternal grand-parent). This is a property of available data from which the pedigree is built. Incompleteness can manifest in two ways: i) when genealogical links between pairs of individuals (i.e. parent-offspring) are missing at random,

i.e. independently of other missing links in the pedigree, which lead to a balanced pedigree, i.e. relative to missing ancestral links along maternal vs. paternal sides, but an incomplete pedigree, or ii) when links are missing not at random given that there could be individual heterogeneity in the probability of being included in the dataset or not, giving an asymmetrical incomplete pedigree (see Figure S1 for an illustration). For example, among-individual genetic differences in dispersal propensity could result in the underrepresentation of more dispersive individuals in the reconstructed pedigree.

### Study system

For a number of modern human populations, historical data on births, deaths, and marriages is available from civil or parish registers. This information has been used to reconstruct genealogies that were subsequently exploited in QG studies (e.g., Blomquist 2019; see Bolund et al. 2016 for a review and Stearns et al. 2010 for a list of medical-and clinical-based multigenerational datasets). The pedigrees used in this study were reconstructed from the history of Catholic individuals and families who lived in Québec, Canada, between the foundation of New France by French settlers, in 1608, and 1960. Marriage, baptism and burial acts recorded in parish and civil registers have been compiled and linked since the 1970s in two large databanks: BALSAC (http://balsac.uqac.ca/; Bouchard 1989; Vézina & Bournival 2020) and the Ancient Québec Population Register (AQPR, Dillon et al. 2018). Altogether they contain all Catholic marriages celebrated during this period (first marriage in 1621), as well as births and deaths of all individuals for some regions and periods. For this study, we used data from BALSAC, which also contains part of AQPR data, and now comprises 4 million digitized records of events representing the lives of 5 million individuals. Those dating from 1621 are duplicate copies of church registers, while those dating from 1900 onwards are from civil registers.

This dataset does not represent a homogenous population but rather encapsulates the colonization history of the Québec territory, which took place in multiple waves, in different regions in and time periods. BALSAC data include the locality of records, i.e., parishes or municipalities, which are grouped into 23 regions (Figure S2; see http://balsac.uqac.ca/ for more details). Pedigrees for these regions vary in their attributes and quality. We selected six of them to assess how this variation impacted QG analyses (see SI for selection criteria): Bas-St-Laurent, Bois-Francs, Charlevoix, Côte-de-Beaupré, Gaspésie, Laurentides. Altogether, these regions comprise 5% of individuals in BALSAC, or ∼250,000 individuals in total. Table 2 provides quantitative descriptors of each pedigree.

**Table 2.** Summary pedigree statistics and genealogical properties of the final six pedigrees pruned for quantitative genetic analyses.

|  | Gaspésie | Bas-St-Laurent | Laurentides | Charlevoix | Côte-de-Beaupré | Bois-Francs |
| --- | --- | --- | --- | --- | --- | --- |
| **Records** | 1,949 | 3,741 | 5,055 | 6,878 | 8,450 | 12,032 |
| **Maternities** | 1,288 | 2,473 | 3,298 | 6,074 | 7,057 | 9,071 |
| **Paternities** | 1,288 | 2,473 | 3,298 | 6,074 | 7,057 | 9,071 |
| **Full sibs** | 1,755 | 4,054 | 4,213 | 12,145 | 9,543 | 14,281 |
| **Maternal sibs** | 1,800 | 4,138 | 4,309 | 12,517 | 9,892 | 14,664 |
| **Paternal sibs** | 1,839 | 4,279 | 4,559 | 13,512 | 10,307 | 15,330 |
| **Maternal grandmothers** | 446 | 722 | 1,218 | 4,649 | 5,025 | 4,773 |
| **Maternal grandfathers** | 446 | 722 | 1,218 | 4,649 | 5,025 | 4,773 |
| **Paternal grandmothers** | 310 | 605 | 1,127 | 4,463 | 5,159 | 4,654 |
| **Paternal grandfathers** | 310 | 605 | 1,127 | 4,463 | 5,159 | 4,654 |
| **Founders** | 661 | 1,268 | 1,757 | 804 | 1,393 | 2,961 |
| **Max. pedigree depth** | 4 | 6 | 6 | 8 | 9 | 7 |
| **Mean maternal sibship size** | 2.69 | 2.84 | 2.46 | 3.65 | 2.72 | 2.90 |
| **Mean paternal sibship size** | 2.71 | 2.9 | 2.57 | 3.83 | 2.80 | 3.01 |
| **Mean inbreeding coefficient** | 0 | $5.42 \times 10^{-5}$ | $6.44 \times 10^{-6}$ | $2.48 \times 10^{-3}$ | $9.36 \times 10^{-4}$ | $8.68 \times 10^{-5}$ |
| **Mean completeness** | 58.88 | 39.03 | 41.45 | 48.10 | 43.61 | 40.65 |
| **Mean expected entropy** | 1.36 | 1.34 | 1.49 | 2.85 | 2.92 | 1.85 |
| **Var. in expected entropy** | 0.29 | 0.48 | 0.57 | 1.87 | 2.48 | 0.86 |
| **Mean rate of missing links** | 0.94 | 0.87 | 0.86 | 0.58 | 0.71 | 0.77 |
| **Mean observed entropy** | 1.26 | 1.05 | 1.08 | 1.00 | 1.31 | 1.07 |

### Missing links and genealogical error rates

Missing parentages and the occurrence of some genealogical errors are hardly avoidable when pedigrees for entire human populations are reconstructed from historical civil and church registers (typically by linking acts such as baptisms, marriages, and burials). Over-linkage refers to incorrect genealogical links made between individuals (i.e., wrong biological parental assignments). They can result from hidden adoptions by social (non-biological) parents or false parentage declarations, for example. Under-linkage refers to missing parental assignments due to information that is lacking or unclear on historical acts. For more information on linkage and data quality, we refer the reader to internal BALSAC documents available at https://balsac.uqac.ca/bibliographie-selective/, in addition to published works such as Vézina & Bournival (2020) and Vézina et al. (2018).

We calculated the degree of missing parentage for each pedigree analyzed in this study as the number of individuals with no known parents over the total number of individual records. The range of missing parentage extends from 5% to 31% per pedigree. Missing parentage is partly due to population founders, whose parents are, by definition, from the ancestral (source) population and thus absent from the pedigrees. Indeed, the pedigree with the highest missing parentage has the largest number of founders. Exclude those, the degree of missing parentage drops dramatically in most of the pedigrees. This means that most non-founders were assigned parents (whether correctly or not). Additionally, pedigree records of paternities and maternities (including grandparentage) are equal within each pedigree, meaning there is no imbalance in terms of missing parentage assignments (Table 2).

Using haplogroup genetic data, the genealogical error rate related to over-linking for BALSAC data was estimated to be 0.55% in maternal genealogical lineages and 0.82% in paternal genealogical lineages (Jomphe 2011). This was confirmed recently by Doyon et al. (*in prep*.), who analyzed mitochondrial DNA sequences (matrilineal inheritance) and Y-chromosome markers (patrilineal inheritance) from living individuals connected to the Québec genealogy. From gene dropping simulations they estimated that 0.80% (s.e.=0.07%) of father-offspring links were erroneous, which includes 0.41% of extra-pair paternities (EPP). As for mother-offspring links, the error rate was estimated to be 0.383% (s.e.=0.009%). Since a pedigree necessarily contains equal numbers of mother-and father-offspring links, the total error rate is thus expected to be ∼0.6%, hence one for every ∼166 links. Note that a mismatch between parent’s and offspring’s DNA needs not reflect a linkage error but could be due to a mutation. Therefore, the per-marker mutation rate was also accounted for in the estimation of the above genealogical error rates. We used the above rates in the present study to check the sensitivity of the results to these parameter values. We also ran the same set of analyses assuming a 1% error rate for both parental lines, i.e., nearly twice the overall rate estimated by Doyon et al. (see SI for these results).

### Pedigree simulations

The pedigrees analyzed in this study are ‘observed’ pedigrees that potentially contain non-negligible amounts of errors. Using the observed pedigree we simulate a plausible ‘true’ pedigree, based *a priori* information on rates and types of genealogical errors. Phenotypic and breeding values for a Gaussian trait were simulated across the simulated ‘true’ pedigree. To assess the impact of pedigree attributes, we compared genetic parameters estimated from the observed pedigree to the preset parameters used to simulate breeding values using the hypothetical ‘true’ pedigree. Note that the term ‘true’ refers to the true pedigree structure relative to the phenotypic data that are used in power and sensitivity analysis and does not necessarily reflect a pedigree that actually exists in nature (Morrissey et al. 2007). The ‘true’ pedigree then is completely based on the observed pedigree and the only difference between them is that the former is considered to contain no errors and is based on the true breeding structure of the population under consideration (Morrissey & Wilson 2010).

The specific analysis framework is the ‘reverse’ approach, as implemented in the R package *pedantics* (Morrissey & Wilson 2010). We simulated plausible ‘true’ pedigrees for each of the six regions using the rpederr function, which “probabilistically assigns “true” parents given an incomplete and potentially erroneous pedigree” (Morrissey 2018). To use this function, we re-arranged the pedigree dataset as follows. First, we filtered out maternal and paternal lineages with less than 10 individuals. Such small lineages result either from a true lineage end (i.e., no more descendants) or from the fact that subsequent descendants were born outside the focal region (i.e., emigration). Without this filtering of small lineages, pedigrees would be quite imbalanced in some cases. In the case of both maternal and paternal lineages, the average proportion of lineages < 10 individuals make up only 11% of total lineage sizes across the studies pedigrees. Each known social parent-offspring dyad remaining in the filtered dataset was assigned a probability of error corresponding to the genetically-estimated genealogical error rate (i.e., 0.0080 and 0.00383 for father-and mother-offspring links, respectively). If the identity of the mother or father was missing, the probability of erroneous parental assignment was set to 1. Importantly, this does not mean that the parental assignment is assumed to be erroneous with 100% certainty. Rather, setting this value to 1 indicates to the function that the parental assignment is missing or null (in the case of a founder) for these individuals, thus allowing their inclusion in the process of simulating plausible ‘true’ pedigrees. Founders or immigrants arriving later (i.e. post-foundation) are individuals with no record of birth year and parent IDs, hence were also assigned an erroneous parental assignment probability of 1 (we confirmed with BALSAC staff the founder/immigrant status of these individuals). Finally, the sex, birth year, and cohort of first and last offspring known for each individual were included as required input variables to identify the pool of potential parents for individuals with either missing or (simulated) erroneous parentage. The rearranged pedigree data was fed to the rpederr function, which provides as output a plausible ‘true’ pedigree. The latter then served as input in the next step, namely the simulation of individual phenotypic and breeding values.

Since our aim was to assess how genealogical attributes affect the reliability of QG estimates, we calculated the following pedigree metrics for the six datasets: mean lineage completeness (Cazes & Cazes 1996), average inbreeding coefficient (Thompson 1986), total number of founders, and mean entropy (based on Kouladjian 1986, also see Cazes & Cazes 1996 and Milot et al. 2011). These metrics were obtained from the *GENLIB* R package (Gauvin et al. 2015; see SI for how metrics are calculated).

### The animal model

The animal model is a mixed-effect linear regression which decomposes phenotypic trait variation into its genetic and non-genetic components. The individual’s breeding value is the deviation of the phenotypic value from the population average that is explained by the additive effects of the individual’s genes. In its simplest form, the phenotype *z* of an individual *i* can be written as: *z*_*i*_ = *µ* + *a*_*i*_ + *e*_*i*_, where *µ* is the population mean, *a*_*i*_ is the additive genetic value for individual *i*, and *e*_*i*_ is the residual error. Both the additive genetic effect and residual error are assumed to be normally distributed. The model also accommodates other sources of variation to be fitted as random effects, such as maternal or shared familial environment effects. In statistical terms, the pedigree provides the *n* × *n* matrix **A** of the genetic relatedness between every pair made of the *n* individuals in the population. The animal model uses the inverse (**A**^-1^) of this matrix in combination with individual phenotypic data to estimate QG parameters.

Parametrization of the animal model can have a considerable effect on the reliability of QG estimates (Kruuk & Hadfield 2007; Wilson 2008; Wolak et al. 2015). For example, certain phenotypic traits are not normally-distributed, or may be skewed towards a given value (e.g., zero-inflated distributions), hence proper parametrization is crucial (de Villemereuil 2018), especially for fitness (Bonnet et al. 2019). Given that phenotypic variation is most often due to more than one source, it is important to specify the variance structure model adequately, otherwise some variance terms can be poorly estimated (e.g., inflated when some important variance sources are missing; Kruuk and Hadfield 2007).

Pedigree individuals who are informative about the genetics of quantitative traits are those either with measured phenotypes and genetically related to one or more other phenotyped individuals, or ancestors who contribute to make a genealogical loop between phenotyped individuals. Computational algorithms to fit QG models are more efficient when uninformative individuals are removed from the pedigree. Therefore, we pruned pedigrees to keep only the informative individuals, using the `prunePed` function in the *MCMCglmm* R package (Hadfield 2010). Additionally, the phenotypic dataset supplied to the model was pruned to retain only individuals for which the year of birth is known and had at least one successful reproductive event. The reasoning behind this is that, at the least, only individuals with a known year of birth and reproductive event can be phenotyped by having their life history reconstructed. Furthermore, this decision allowed us to be conservative by checking for the reliability of estimates based on a restricted phenotypic dataset reflective of the case of QG analysis in natural populations. We used the Bayesian implementation of the animal model in the *MCMCglmm* R package employing Markov chain Monte Carlo methods (Hadfield 2010) in the R environment (v3.6.3; R Development Core Team 2019). All models were run for 4,000,000 iterations after an initial burn-in of 500,000 iterations and sampled every 4,000 iterations (i.e., 1,000 samples kept) using uninformative priors.

### Simulation analyses

We used the phensim function from *pedantics* to simulate phenotypic values for each individual in the simulated ‘true’ pedigree. Phenotypic values were calculated as the sum of additive genetic effects (i.e. breeding values) and environmental effects simulated from user-specified (co)variance matrices. We ran scenarios with one and two response traits, in the latter case to assess the impact of pedigree quality on genetic covariance estimation. Additionally, in one scenario, we simulated maternal environmental effects to replicate phenotypic resemblance due to the sharing by siblings of the familial environment. In total, we ran three scenarios, as follows:

- **Scenario 1**: Univariate response with simple trait architecture: *z*_*i*_ = *µ* + *a*_*i*_ + *e*_*i*_, where *µ* is the population mean (with *µ* = 0 in our simulations), *a*_*i*_ is the additive genetic value for individual *i*, and *e*_*i*_ is the residual error. The phenotypic variance is *V*_*P*_ = *V*_*A*_ + *V*_*E*_ = *1*, where *V*_*A*_ is the additive genetic variance and *V*_*E*_ is the residual (environmental) variance. *V*_*A*_ was set to 0.1 or 0.3, with *V*_*E*_ = 1-*V*_*A*_.
- **Scenario 2**: Univariate response as in Scenario 1 but adding the term *m*_*i*_ for shared familial environment effects: *z*_*i*_ = *µ* + *a*_*i*_ + *m*_*i*_ + *e*_*i*_. Here, *V*_*P*_ = *V*_*A*_ + *V*_*M*_ + *V*_*E*_ = *1. V*_*A*_ was set to 0.1 or 0.3, *V*_*M*_ was set to 0.1, with *V*_*E*_ = 1-(*V*_*A*_ + *V*_*M*_).
- **Scenario 3**: Bivariate response with traits *z*_*1*_ and *z*_*2*_: *z*_*i*_ = *µ* + *a*_*i*_ + *e*_*i*_. Again, *V*_*P*_ = *V*_*A*_ + *V*_*E*_ = *1. V*_*A*_ was set to 0.1 or 0.3 for trait *z*_*1*_ and fixed to 0.1 for *z*_*2*_. The genetic correlation *r*_*g*_ between *z*_*1*_ and *z*_*2*_ was set to 0.1 or 0.5, to simulate a weak and strong correlation, respectively.

Under the first two scenarios, we ran five simulations for each regional pedigree × *V*_*A*_ value, for a total of 10 simulations. Due to computational time constraints (several weeks for a single run of a bivariate model) when fitting bivariate models, we ran a single simulation per pedigree and parameter value in the case of Scenario 3.

### Reliability measures

For Scenarios 1 and 2, precision and accuracy for *V*_*A*_ and *V*_*M*_ estimates were first checked visually by plotting posterior modes as point estimates and 95% highest posterior density (HPD) intervals as precision estimates from posterior distributions of parameters. For each parameter estimated in the models, we calculated the mean squared error as: 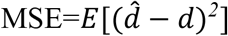, where *d* is the true parameter value (i.e., the value specified to simulate phenotypes) and 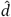 the value estimated from the animal model fitted to simulated trait data. The root mean squared error (RMSE) was then used to standardize the scale of estimates from the different models in order to compare them (de Villemereuil et al., 2013). For a given simulation, the posterior distribution of the RMSE for a given parameter was provided by the MCMC sample of 1,000 iterations by applying the same calculation over every saved MCMC iteration. The final results shown here are based on the average of the five simulations for Scenario 1 and Scenario 2).

### Interpretation of posterior distributions

The 95% HPD interval provides a measure of the uncertainty about the true parameter value (they are not to be confused with confidence intervals). This means the larger the HPD interval is the wider the range of parameter values that are plausible (i.e., a lower precision). We also used two Bayesian indices to draw inferences from the posterior distribution about the existence and significance of a given genetic or environmental effect on a trait (see Makowski et al. 2019b). The Region of Practical Equivalence (or ROPE)-based index, specifically the *ROPE* (95%), corresponds to the proportion of the 95% HPD that lies within the ROPE (loosely speaking, this a Bayesian version of the frequentist’s concept of “power”). This type of index involves redefining the alternative hypothesis from the classic point-null (i.e., the value 0) to a range of values considered negligible or too small to be considered practically relevant to the biological question being studied (Kruschke 2014). According to Makowski et al. (2019b), the ROPE-based index offers information in favor of the null hypothesis and is sensitive to sample size (see SI for information on how the *ROPE* range was defined in our study). We also used the *Probability of Direction* (*pd*) which is the Bayesian equivalent of the frequentist *p*-value in that it reflects the existence of an effect. This index varies between 50% and 100%, a high value suggests that an effect exists while a low value indicates uncertainty around its existence. To calculate these indices in our study, we used the R package *bayestestR* (Makowski et al. 2019a).

## Results

### Pedigrees

Following pedigree reconstruction and filtering, we ended up with six datasets corresponding to six Québec regions. There was a large amount of variation between the datasets in terms of structure and properties (Tables 2 and 3). This variation is especially obvious when we contrast the pedigrees of the Gaspésie and Bois-Francs regions. The sample size and number of founders are roughly five times larger in the Bois-Francs than the Gaspésie pedigree; pedigree depth is 4 for Gaspésie and 7 for Bois-Francs. Yet, these two pedigrees have some similarities, such as the mean inbreeding coefficient and expected genealogical depth (Tables 2 and 3, Figures S6 and S7).

### Power

Simulated values of *V*_*A*_ or *r*_*g*_ are highly correlated (*r* > 0.99) with the realized value within each simulated population. Barring one exception, all pedigrees across scenarios allowed the detection of the existence of the genetic component in traits’ architecture. Only Gaspésie had a reduced power to detect the genetic signal in the simulated trait according to the *ROPE* index (Figure 6).

### Precision and accuracy

Visual inspection of the posterior distributions (Figures 1-5) reveals that larger and deeper pedigrees provide a higher precision in *V*_*A*_ and *Cov*_*A*_ estimates but that their accuracy depends on trait architecture (i.e., with vs. without the shared familial environment effect) and model parameterization. This is congruent with differences in RMSE among pedigrees and across simulation scenarios (Table S1). Under Scenario 1, larger and deeper pedigrees (Bois-Francs, Côte-de-Beaupré) resulted in the lowest RMSE when *V*_*A*_ was set to 0.1 (0.011 [0 - 0.037] and 0.005 [0 - 0.035], respectively). When *V*_*A*_ was set to 0.3, the pedigree of the Laurentides region gave a slightly better accuracy, albeit a lower precision (larger HPDs; 0.008 [0 - 0.080]), than the much larger one for Bois-Francs (0.012 [0 - 0.059]; Table S1). For Scenario 3, which involved a single simulation per pedigree and per parameter, a visual inspection was done by directly plotting the posterior distribution using the R package *bayesplot* (Gabry et al. 2019).

**Figure 1.**
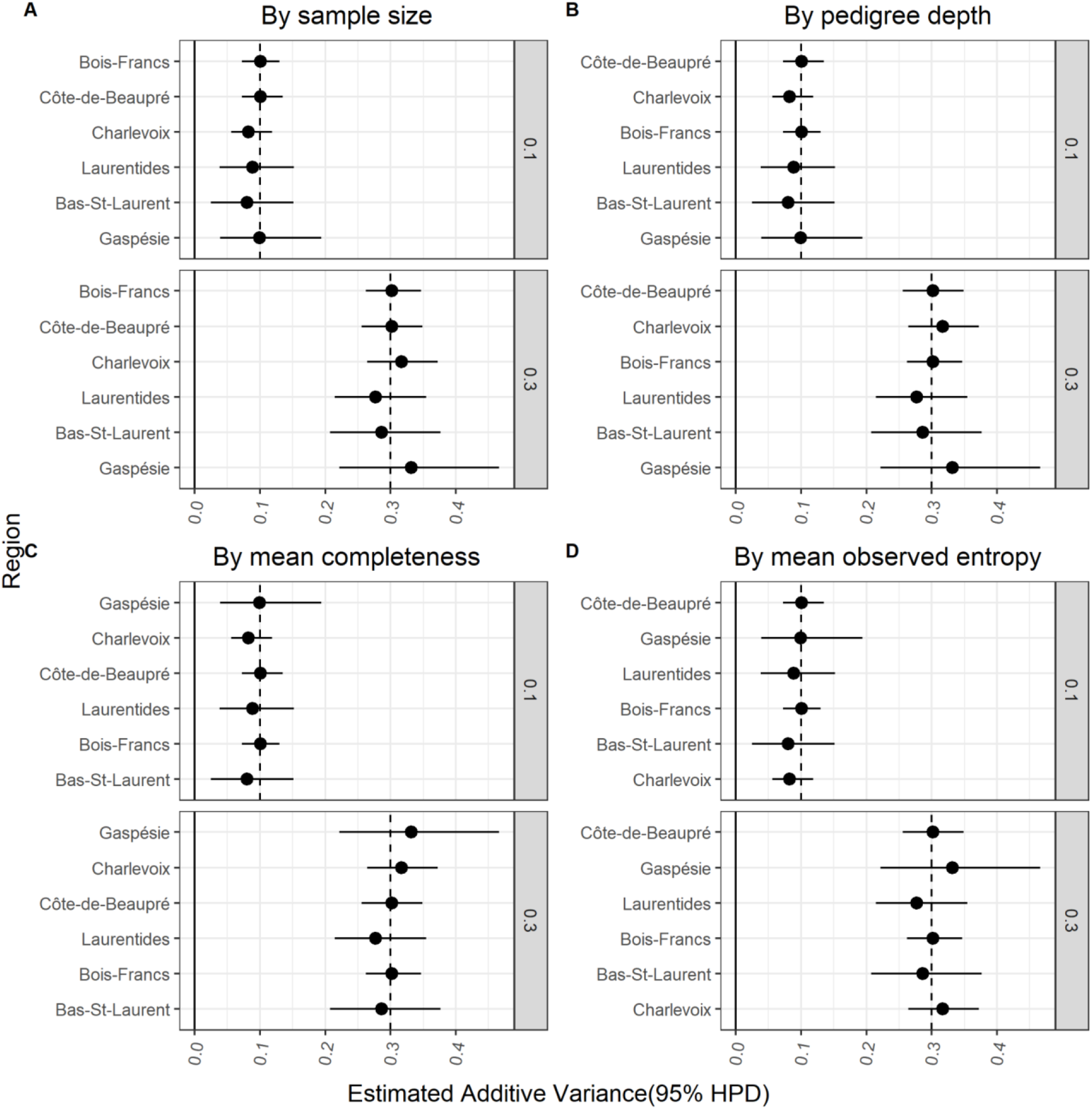
Estimates of additive genetic variance for each of the six pedigrees based on Scenario 1 (univariate simple phenotype; *V*_*A*_ = *0*.*1 and 0*.*3*). In each plot, pedigrees are sorted, from bottom to top, by increasing: **A**. sample size, **B**. pedigree depth, **C**. mean completeness, and **D**. mean observed entropy. Each dot represents the average posterior mode-estimates, along with the average 95% HPD intervals based on the five simulations per pedigree per parameter value. Vertical dashed lines represent the expected value of the variance.

When another source of resemblance between relatives was added (i.e., shared familial environment, Scenario 2), precision and accuracy decreased (average RMSE; Figure 2). When both *V*_*A*_ and *V*_*M*_ were set to 0.1, the pedigree of the Laurentides region had the lowest RMSE (0.048 [0.009 - 0.139]). In the case where *V*_*A*_ was set to 0.3, the pedigree of the Charlevoix region gave the best accuracy and precision compared to the other pedigrees (0.028 [0 - 0.089]). On average, *V*_*A*_ was overestimated by 46% and *V*_*M*_ underestimated by 42%. Even larger and deeper pedigrees produced inaccurate *V*_*A*_ and *V*_*M*_ estimates, to the extent that that in some cases, the average HPDs of the posterior distribution of *V*_*A*_ over five simulations did not even contain the expected value. RMSEs show that larger and deeper pedigrees were no longer performing better than the smaller ones *V*_*M*_ was fitted. We redid the analysis with a less restrictive phenotypic datasets by keeping individuals without a known year of birth which increased the mean maternal family sibship size to ∼6 across pedigrees. Results from this analysis show the same bias in *V*_*A*_ and *V*_*M*_ estimates (not shown).

**Figure 2.**
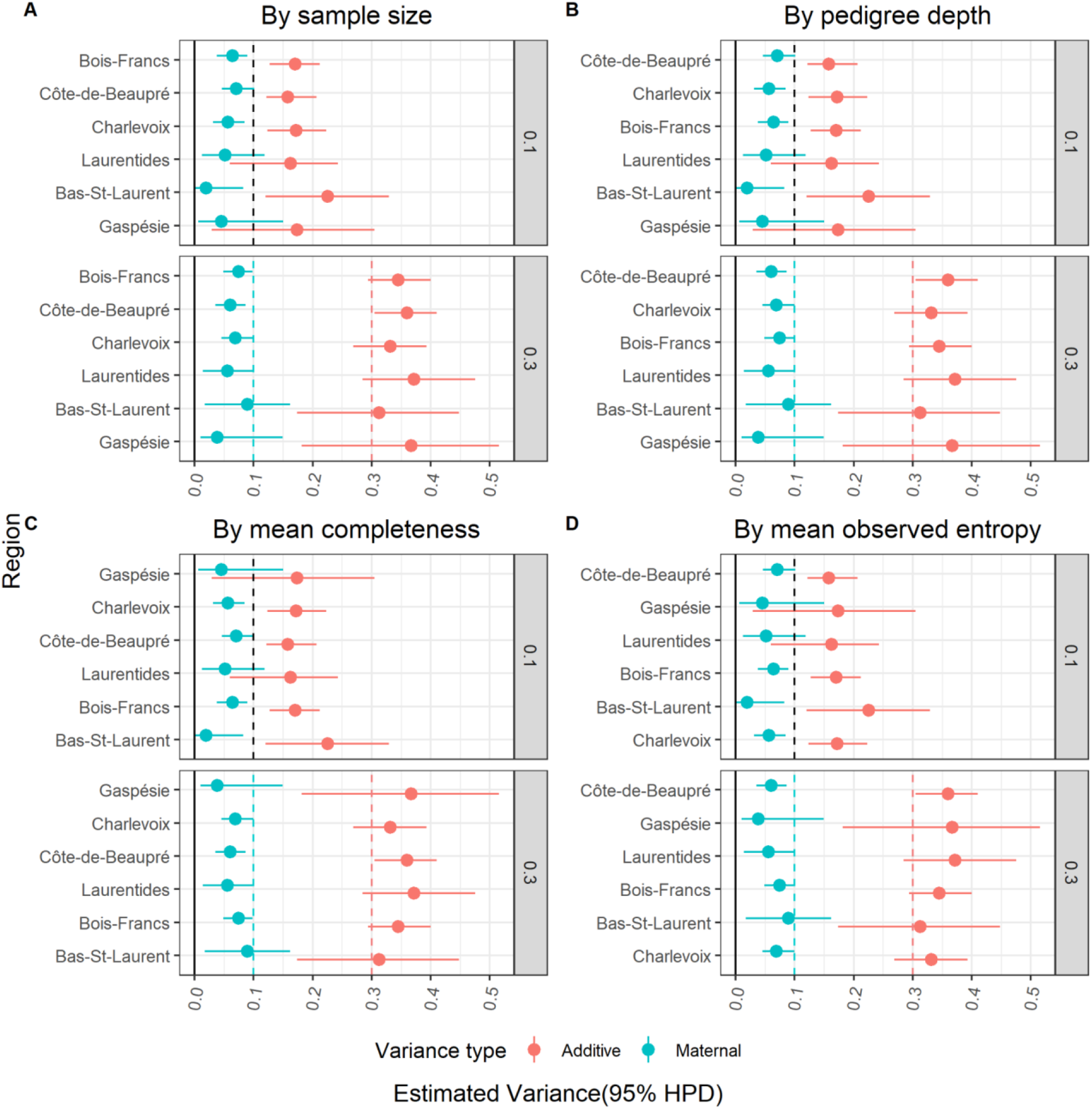
Estimates of additive genetic variance and maternal variance for each of the six pedigrees based on Scenario 2 (univariate complex phenotype; *V*_*A*_ = *0*.*1 and 0*.*3*;*V*_*M*_ = *0*.*1*). In each plot, pedigrees are sorted, from bottom to top, by increasing: **A**. sample size, **B**. pedigree depth, **C**. mean completeness, and **D**. mean observed entropy. Each dot represents the average posterior mode- estimates, along with the average 95% HPD intervals based on the five simulations per pedigree per parameter value. Vertical dashed lines represent the expected value for each variance component.

**Figure 3.**
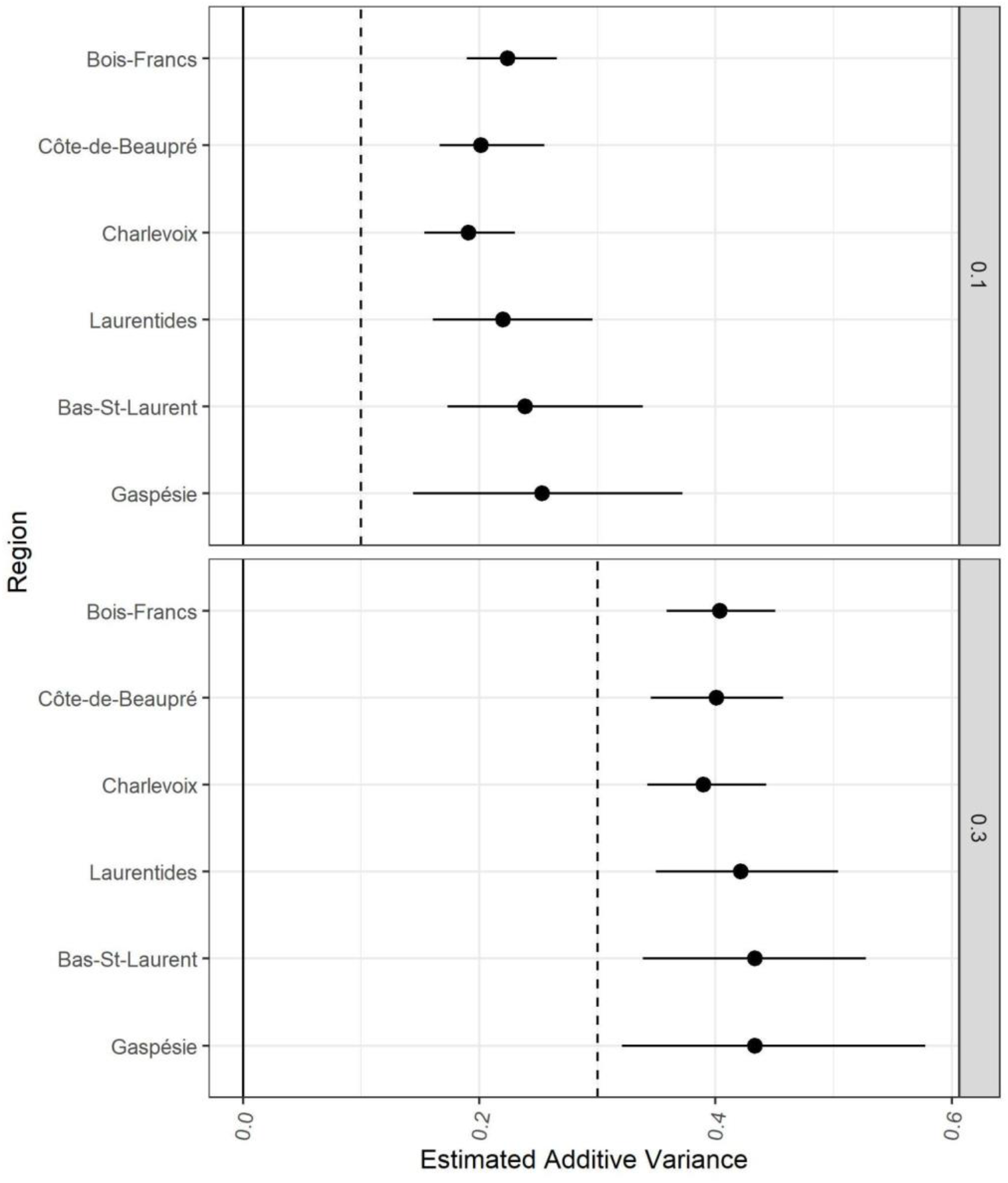
Estimates of additive genetic variance for each of the six pedigrees based on Scenario 2 with improper parameterization (univariate complex phenotype; *V*_*A*_ = *0*.*1 and 0*.*3*;*V*_*M*_ = *0*.*1*). Each dot represents the average posterior mode-estimates, along with the average 95% credible intervals based on the five simulations per pedigree per parameter value. Vertical dashed lines represent the expected value of the variance

**Figure 4.**
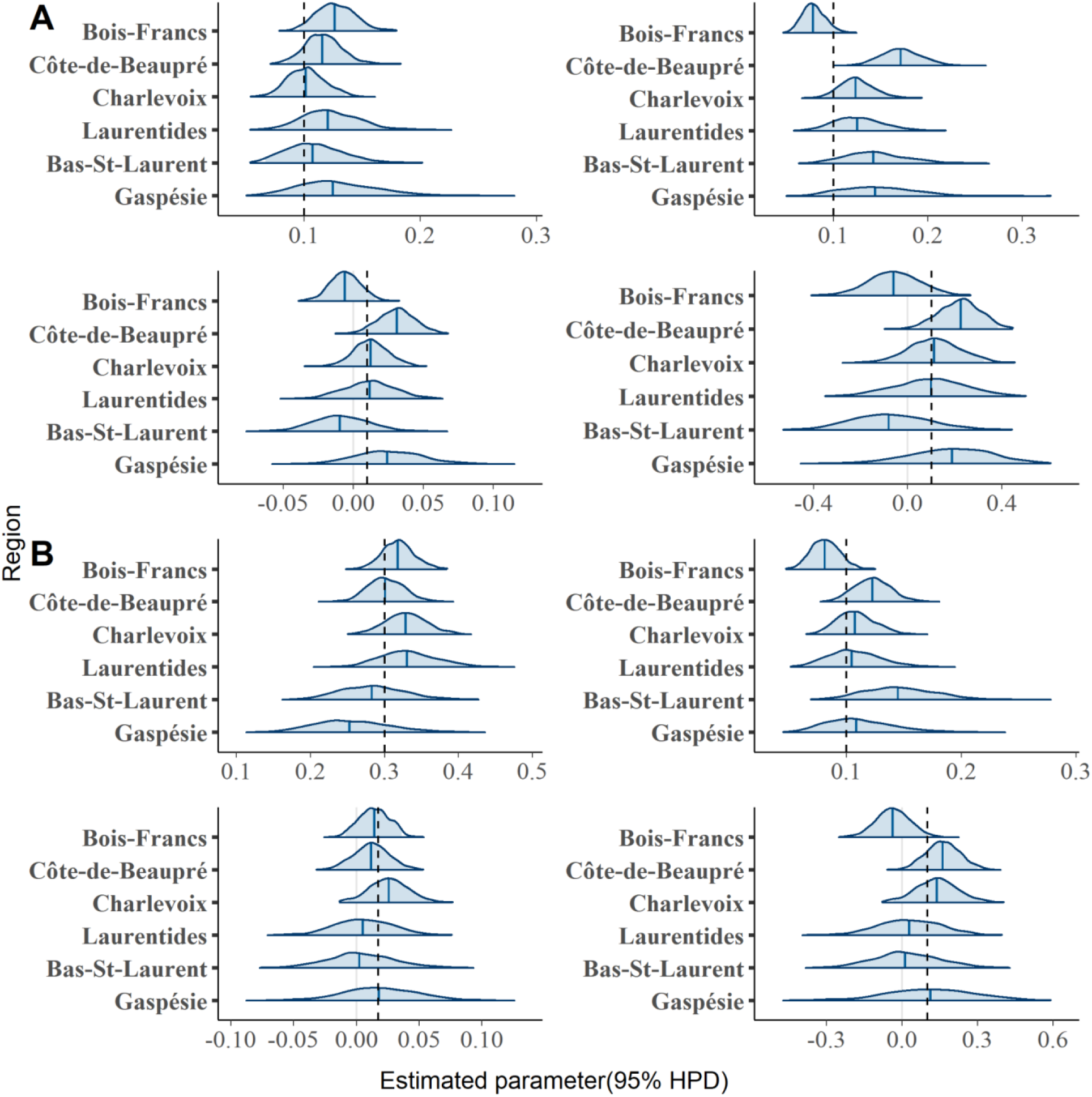
Posterior distributions of estimated quantitative genetics parameters (with the 95% HPD interval) for each of the six pedigrees based on Scenario 3 (bivariate simple phenotype with genetic correlation *r*_*g*_= 0.1). Top left is additive variance of trait *z*_*1*_, top right is additive variance of trait *z*_*2*_, bottom left is additive covariance and bottom right is genetic correlation. **A**) 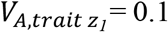 and *Cov*_*A*_ = *0*.*01*; **B**) 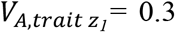 and *Cov*_*A*,_ = *0*.*0173*. In both cases, 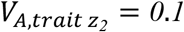. Vertical dashed lines represent the true expected value of the variance.

**Figure 5.**
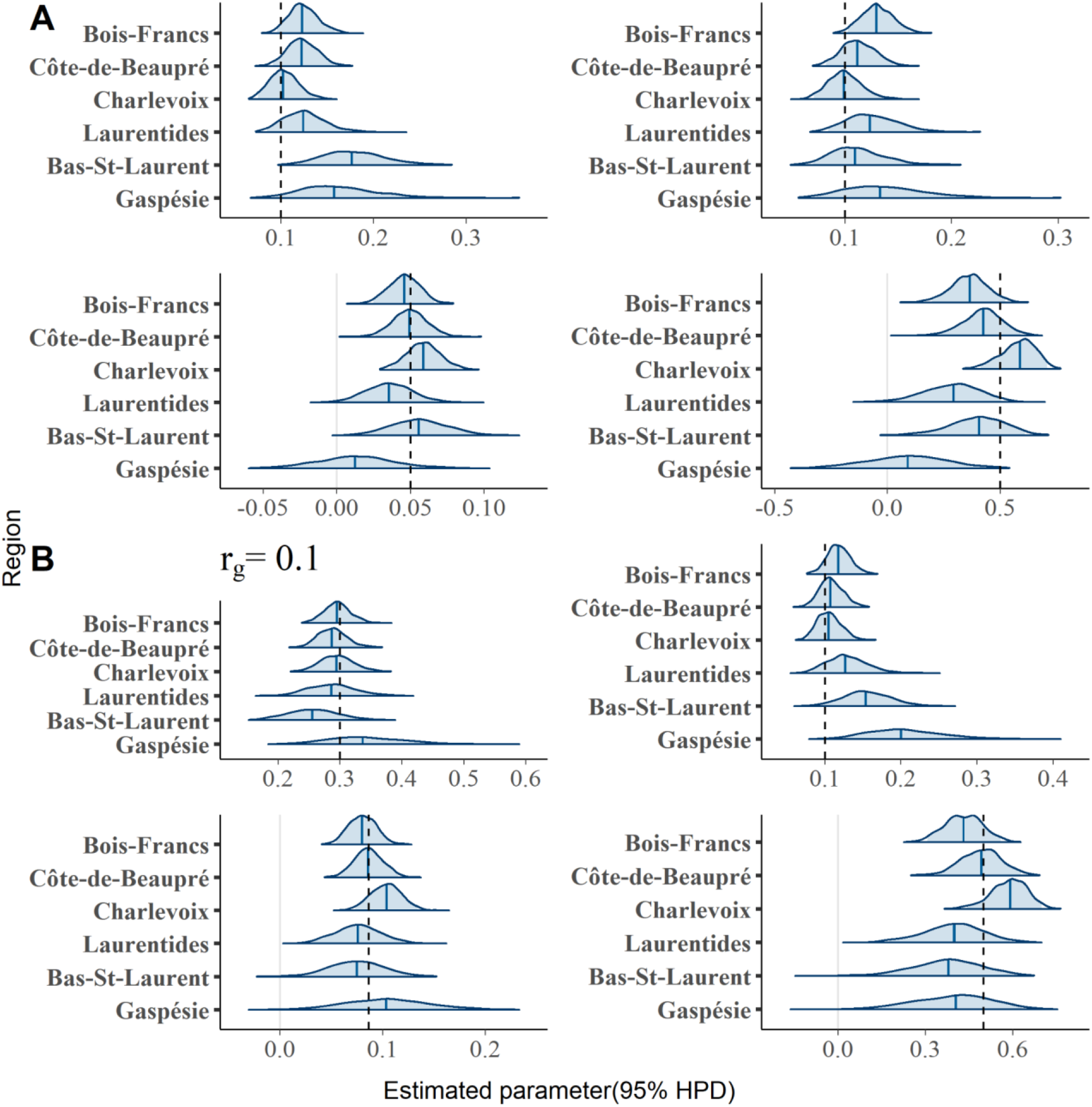
Posterior distributions of estimated quantitative genetics parameters (with the 95% HPD interval) for each of the six pedigrees based on Scenario 3 (bivariate simple phenotype with genetic correlation *r*_*g*_= 0.5). Top left is additive variance of trait *z*_*1*_, top right is additive variance of trait *z*_*2*_, bottom left is additive covariance and bottom right is genetic correlation. **A**) 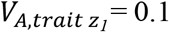 and *Cov*_*A*_ = *0*.*05*; **B**) 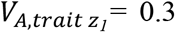 and *Cov*_*A*,_ = *0*.*0866*. In both cases, 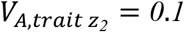. Vertical dashed lines represent the true expected value of the variance.

**Figure 6.**
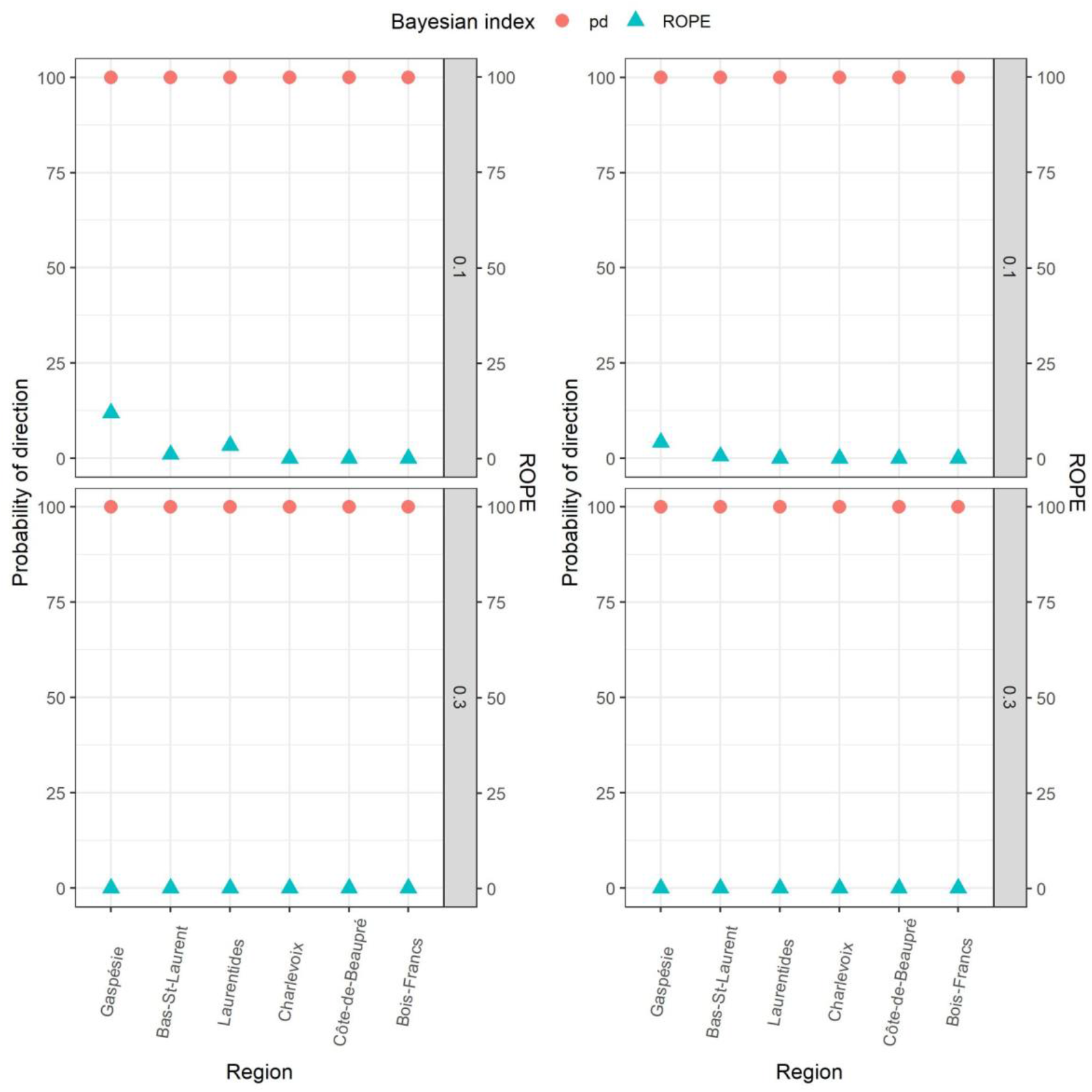
Estimates of Bayesian indices, posterior of direction (*pd*) and *ROPE*, for each of the six pedigrees based on five simulations. *Left*, Scenario 1 (univariate simple phenotype); *Right*, Scenario 2 (univariate complex phenotype). Dots represent the average *pd*; Triangles represent the average *ROPE*.

To assess parameter identifiability under Scenario 2, we measured Spearman’s rank correlation between the posterior distributions of *V*_*A*_ and *V*_*M*_ for each simulation. The average correlation between the two the parameters across all pedigrees was -0.51, indicating a reduced identifiability when both *V*_*A*_ and *V*_*M*_ were fitted together (also confirmed by MCMC autocorrelation checks). Identifiability was slightly higher with larger and deeper pedigrees, dropping from an average correlation of -0.66 for the pedigree of the Gaspésie region to -0.36 for the pedigree of the Charlevoix region.

## Discussion

Assessing a population’s evolutionary potential or change requires reliable estimates of genetic parameters. However, the validity of applying QG analyses to the pedigree reconstructed for a natural population, whether from field observations or genetic markers, has not been systematically investigated with quantitative tools in empirical studies (but see Bourret & Garant 2017), including in some previous work by the authors of this paper. The sensitivity of parameter estimation to pedigree attributes has been explored mostly in (passerine) birds and (ungulate) mammals. This applies to the present study on a mammalian species. Therefore, extrapolation of system-specific conclusions should be done with caution, as other taxa may be characterized by on average different pedigree topologies or tractability (Pemberton 2008). It should not save one from performing sensitivity analysis for the empirical under investigation, when needed. Rather, aforementioned studies can help interpret QG results, design sampling schemes, or foresee the limits the empirical pedigree at hand may have to test genetic architecture models of varying complexity. Overall, our simulation analyses show that all pedigrees tested herein convey enough information to detect and quantify small *V*_*A*_ values with modest uncertainty. Results for *V*_*M*_ and genetic covariances vary more. Below we discuss how pedigree attributes and errors affect the reliability of QG estimation in our population system, and issues linked to model parameterization and identifiability.

### Validity and pedigree error rate

Using a genetically-based estimate of the genealogical error rate for the FC datasets, our study suggests that the reliability of QG estimates is little impacted by such errors. In fact, for a phenotype with simple architecture (Scenario 1), we did not detect a downward bias in the additive genetic variance. However, increasing the error rate to 1%, the upper bound of the expected genealogical error rate in the FC datasets, lead to a slight underestimation and reduction in accuracy (see Figure S3). In a previous sensitivity study, Charmantier & Réale (2005) found that a paternity misassignment rate of 20% is the cutoff where the error rate begins to affect QG estimates. Generally, EPP rates in human populations are considered to be low (range: 0.4‒11.8; Andreson 2006; also see Larmuseau et al. 2016) with a median of 1.7%. However, our results show only a slight increase in the genealogical error rate (i.e., from 0.38% to 1%) is enough to negatively impact QG estimates. We observed this underestimation uniformly across all analyzed pedigrees which rules out the possibility that some of our pedigrees are more sensitive (in terms of QG analysis) to slight increases in the error rate. This makes it more pertinent to investigate if certain particular attributes explain the observed discrepancy when we increase the error rate and avoid generalizing results from studies using pedigrees from wild passerine and ungulate populations to social human pedigrees.

In *pedantics* phenotypic simulations, a trait value can be generated for each individual in the pedigree. However, in empirical studies, phenotypic data is often unavailable for a proportion of these individuals. In our analyses, the final datasets supplied to the animal model reflected the realistic expectation. Each of the six phenotypic datasets were pruned to include only individuals with both a known year of birth and at least one reproductive event. Without this information it would be practically impossible to reconstruct individual life history, a common phenotypic trait of interest in natural populations. Additionally, compared to a compilation of QG studies on wild populations by Postma (2014), the sample size (total number of records) of our pruned phenotypic datasets was considerably higher than the median number of records with life history trait data in these studies ‒ which was 377 ‒ and, in most cases, higher than the upper bound of the range (range 6–4992 records; from 39 studies on 19 species). Sample size and depth are two important factors that determine pedigree quality (Wilson et al. 2010). Accordingly, our results demonstrate that the larger and deeper the pedigree was the more precise the QG estimates were. In fact, both sample size and pedigree depth exhibited the strongest correlation with a pedigree’s RMSE score (Figure S4), however it is difficult to separate their independent effect since they are strongly correlated with each other across datasets (*r* = 0.73). We also found that mean completeness and genealogical entropy exhibited a considerable correlation with RMSE score. It is not completely clear in what manner these two pedigree attributes impact the precision and accuracy of QG estimates. Nonetheless, they should not be ignored in pedigree-based studies testing the validity of their QG estimates based on their respective pedigree.

### Impact of additional source of resemblance

We observe biases in parameter estimates when an additional source of phenotypic similarity between relatives is included in the model (Scenario 2). Though larger and deeper pedigrees still provided fairly precise estimates, only those from pedigrees with a high mean completeness also exhibited good accuracy. This is a problem because life history traits are known to exhibit non-genetic sources of resemblance between relatives (e.g., Laugen et al. 2002, Pettay et al. 2005). In addition, when genetic parameters are estimated to test evolutionary hypotheses, accuracy is more important than precision because overestimation (or underestimation), even with low uncertainty, will lead to faulty interpretations. We can think of one reason that could explain the overestimation of the additive genetic variance observed under Scenario 2. The proportion of individuals with less than two offspring was relatively high in pruned pedigrees, which could have negatively impacted the power to estimate shared familial environment effects. However, redoing the analysis using less restrictive phenotypic datasets thus increasing the mean number of siblings per mother included in the pedigree did not eliminate the estimation bias (Figure S5).

### Proper parameterization and low identifiability

Another important factor affecting the precision of estimates, and the validity of QG analyses in general, is proper model parameterization (Morrissey et al. 2007). This means that the model structure includes the appropriate sources of variation in the focal trait, both genetic and non-genetic. Even a pedigree of very high quality could produce inaccurate results if an important variance parameter is missing (Kruuk & Hadfield 2007). A common issue with wild populations is the overestimation of the additive genetic variance when maternal (family) effects are not included in model specification. This arises because the analysis cannot distinguish well (or at all) the genetic and non-genetic contributions to the resemblance between relatives (such as (half)siblings sharing the same mother). When we misspecified the animal model by dropping the maternal variance under Scenario 2, all of the expected maternal variance was rather included into the additive genetic variance. This result was consistent across all pedigrees which is surprising because we would expect that deeper pedigrees would perform better compared to pedigrees with less depth. However, this was not the case which highlights the fact that there is no situation where omitting the shared maternal environment effect (or any other non-genetic source of resemblance), when it is expected to contribute to the trait’s phenotypic variation, does not lead to estimation issues. Additionally, our results revealed a low identifiability between the genetic and shared familial environment components. This is similar to the results found by Bourret & Garant (2017) using a frequentist framework, and here we confirm the same pattern using a Bayesian framework.

### Validity of joint estimates when using bivariate animal models

In evolutionary QG studies, reliable estimates of additive genetic covariances should be on par to that of additive genetic variances. Covariance between a focal trait and fitness (Robertson-Price identity) determine the genetic response to selection, while covariance among multiple traits can set constraints on this response or reflect evolutionary trade-offs of interest. Our Scenario 3 involved bivariate models mimicking such a situation where a life history trait is either strongly or weakly genetically correlated with fitness. Though it was difficult to perform multiple simulations per pedigree given prohibitive computation time, our results point out to conclusions about the reliability covariance estimates similar to those for the additive variance. Namely, bigger and deeper pedigrees provided more precise estimates, but nonetheless inaccurate in some cases. An important factor affecting these estimates was the magnitude of the simulated covariance between the two traits: a smaller simulated covariance resulted in poorer estimates from subsequent animal model analyses. Though in all cases (but one) the most probable sign of the estimated covariance matched that of the simulated parameter. These results reveal that studies interested in genetic covariances and correlations, either in the context of studying genetic trade-offs or the genetic response to selection, should consider the possibility that, regardless of its attributes, certain pedigrees might be underpowered to detect weak genetic covariances.

## Conclusion

High density genomic coverage is becoming increasingly available for natural study systems, allowing genome-wide estimation of the realized relatedness between individuals, as opposed to pedigree-based relatedness expectations. This can definitely improve the estimation of QG parameters, in addition to opening the possibility of studying the QG of a natural system where it is difficult to obtain a multi-generational pedigree. In their review of empirical and simulation studies of QG in wild populations using marker-based estimates of relatedness, Gay et al. (2013) determined that the method of pedigree reconstruction using genetic markers performs poorly in parameter estimation relative to the method of relatedness matrix reconstruction using high density genetic markers (i.e., GRM, or genomic relatedness matrix). The results of our study show that socially-reconstructed pedigrees can still perform remarkably well in terms of QG parameter estimation (ex. Bérénos et al. 2014). The combination of inexpensive sequencing of genomes and increased coverage of genetic information along the pedigree will no doubt better the validity of key genetic parameter estimates.

Though REML (Restricted Maximum Likelihood) is considered a better estimator of additive variance for Gaussian traits, we recommend using a Bayesian approach for simulation studies aiming to assess the validity of estimates for any type of trait distribution for the following reasons: (i) its intuitive interpretation of estimates, (ii) possibility of assigning different degrees of priors, and (iii) checking and controlling for autocorrelation between variance components (but see Houle & Meyer 2015). Though large and deep pedigrees can be *a priori* considered adequate for QG analyses, this comes with the caveats of considering the impact of trait architecture, pedigree quality, and model parameterization. We also highlight in this study the value of investigating the impact of pedigree properties such as completeness and genealogical entropy. With routine sensitivity analyses becoming easier to implement with accompanying packages in the statistical environment R making the programming easier, there is high expectation of producing validity measures alongside reporting QG estimates for either a new pedigree or when testing a more complex model using a previously-studied pedigree.

