## Supplementary information document for "Assessing the impact of pedigree quality on the validity of quantitative genetic parameter estimates"

#### Pedigree filtering criteria

The BALSAC project includes over 4 million records on more than 5 millions individuals over the span of almost four centuries and covering 23 different subregions of French-Canada (Québec, see Figure S5). We separated individual records based on the region of birth and reconstructed the pedigree of each subregion using this information. There was immense variation between the different subregions in terms of sample size, number of generations, number of founders. The major determining factors in quantitative genetic analyses are the size and depth of the pedigree. Hence, our first filtering criteria involved removing the datasets which had a very small sample size < 1,000 individuals and pedigree depth < 4 generations.

Following this, we were left with 13 pedigrees. Our second filtering criteria concerned the long processing and run time for a single model to converge in the case of pedigrees with very large sample sizes. For example, the Saguenay-Lac-St-Jean pedigree with a sample size > 4e+5 required several weeks to run a single iteration of a quantitative genetic model. For this reason, we cut out at least half of the remaining pedigrees exhibiting this issue, which left us with the final 6 pedigrees used in this study. We note that even after these two filtration processes, the variation between the final 6 pedigrees was still large enough to allow for comparison and assessing the impact of differences in pedigree properties on validity (to better visualise this, we provided the visual representation of the pedigree of *Gaspésie*, a small and shallow pedigree, and the pedigree of *Bois-Francs*, a large and deep pedigree, in Figure S7 and S8 respectively).

#### Calculation of genealogical properties

The pedigrees analysed in this study are based on reconstructed genealogies. To assess the impact of variability in the properties and characteristics of these genealogies, we calculated for each of our six pedigrees the following genealogical metrics:

1. *Mean completeness* (Cazes & Cazes 1996): primarily based on the completeness index of a pedigree table $C_x$, i.e. the ratio of the number of known ancestors to the number of expected ancestors, at each generation $x$. Thus, it is calculated using the following formula: $C_x = \frac{number\ of\ known\ ancestors}{number\ of\ expected\ ancestors}$, where the number of expected ancestors at a generation $x$ is equal to $2^x$, the parents' generation being the first one. Finally, we compute the mean value for each pedigree over all the generations.
2. *Average inbreeding coefficient* (Thompson 1986): the inbreeding coefficient of an individual $i$, $f_i$, is the probability that the homologous genes in the two gametes uniting to form individual $i$ are identical-by-descent. Hence, the inbreeding coefficient of an individual is equal to the coefficient of kinship $\psi$ of the individual's parents, i.e. $f_i = \psi(M_i, F_i)$, where $M_i$ and $F_i$ are individual $i$'s parents. Finally, we compute the average inbreeding coefficient over the whole pedigree.
3. *Total number of founders*: in this study, we defined a founder (or a recent immigrant) as an individual missing information on both their parents and their year of birth. We also used the summary statistics from the R package *pedantics* which provides the total

number of founders which matched the raw number calculated from the data. We also had BALSAC run a confirmation analysis on the founders we identified to confirm their founder status in their database.

4. *Expected genealogical entropy* (Cazes & Cazes 1996, based on Kouladjian 1986): this metric is based on an initial metric known as the genealogical entropy introduced by Kouladjian (1986). The three metrics above do not take into account the average number of generations in a pedigree table which makes comparing them between populations (or subpopulations) dubious. To allow comparison, a measure of genealogical depth of a reconstructed pedigree is needed. The measure of entropy in a given genealogy, designating the amount of information contained within it, is calculated by first transforming it into a binary tree where individuals who appear more than once as ancestors are considered as distinct individuals, and then $p_i$ is defined as the probability of the origin of the gene coming from the founder $i$ in this tree. Thus, the entropy parameter $S_B$ is calculated as:

$$S_B = -\sum_i p_i log_2(p_i) = \sum_i N_i \frac{1}{2^{N_i}},$$

where $N_i$ is the generation of founder $i$ and $p_i = \frac{1}{2^{N_i}}$. This parameter gives the expected value of the founders' generation and can serve as a measure of the degree of ancestry. We also calculated the variance in this parameter which is defined as:

$$V = \sum_i p_i {log_2}^2(p_i) - S_B^2 = \sum_i {N_i}^2 \frac{1}{2^{N_i}} - S_B^2,$$

The calculation of the expected genealogical entropy assumes that there are no known missing links. To take into account the impact of missing links, we recalculated the expected genealogical entropy while incorporating an empirical estimate of the rate of missing links:

$$S_B = \sum_{g=1}^{G} N_g \frac{\delta^g}{2^g}$$

where G is the total number of generations over which we are calculating the entropy (g = 1 corresponds to the parent generation, g = 2 the grand-parent generation, etc.). $N_g$ represents the number of ancestors available at generation $g$. $\delta$ represents the rate of missing links, which is the ratio of present genealogical links to the expected genealogical links.

#### Assigning the Region of Practical Equivalence (*ROPE*)

Under the frequentist paradigm it is common practice to test the null hypothesis θ=0, where θ is a population parameter of interest. However, this approach to some extent disconnects "statistical significance" and "effect size" (i.e., biological significance). In this study, we used the *ROPE* Bayesian index to re-interpret 'animal model' results by considering, instead of θ=0, a range of θ values that would be considered as not different enough from zero to be biologically meaningful (see Makowski et al. 2019b). The *ROPE* gives the proportion of a posterior distribution (within 95%-HPD range) that lies below the threshold (i.e., in the range of values considered as not biologically meaningful). While this is based on a subjective choice, this choice is not devoid of scientific criteria, which we make transparent here.

We selected the ROPE range on the basis of the parameter values used when simulating phenotypic values, i.e., $V_A$ was either 0.1 or 0.3. Specifically, having estimates for additive genetic variance ($V_A$) and additive genetic covariance ($Cov_A$) for different datasets, we asked how large should the estimates of these parameters be to be considered biologically meaningful given the initial simulated parameter values. Since what should be considered as "biologically meaningful" is partly subjective, we set different *ROPE*. First, we decided to apply three different ROPE instead of a single one to allow both more conservative (higher ROPE) and more liberal (lower ROPE) interpretations, though in the main text we only report the moderate interpretation. The first one is 5% of the magnitude of the parameter estimate. For example, if simulated $V_A$ was 0.3, then any estimate ≥0.015 would be considered as non-negligeable, which might be viewed as liberal. We also applied a conservative 25% threshold, meaning that a substantial fraction (≥1/4) of a parameter estimate should be within the range to be considered as meaningful. Finally, we also applied a moderate 10% threshold which is the measure reported in the main text.

#### Tables

**Table S1**. RMSE values calculated for each pedigree for each scenario and simulated parameter value. Values are posterior modes of the derived posterior distributions of RMSE, along with 95% highest posterior density (HPD) intervals in brackets. The asterisk corresponds to estimates based on the average of 5 simulations.

| Scenario | Gaspesie | Bas-St-Laurent | Laurentides | Charlevoix | Côte-de-Beaupré | Bois-Francs |
|---|---|---|---|---|---|---|
| Scenario 1 ($V_A$=0.1) | 0.043 [0.009 - 0.118]* | 0.026 [0.002 - 0.076]* | 0.015 [0 - 0.066]* | 0.016 [0 - 0.042]* | 0.005 [0 - 0.035]* | 0.011 [0 - 0.037]* |
| Scenario 1 ($V_A$=0.3) | 0.024 [0 - 0.157]* | 0.039 [0 - 0.115]* | 0.008 [0 - 0.080]* | 0.023 [0 - 0.078]* | 0.015 [0 - 0.058]* | 0.012 [0 - 0.059]* |
| Scenario 2 ($V_A$=0.1) | 0.082 [0.001 - 0.198]* | 0.130 [0.040 - 0.223]* | 0.048 [0.009 - 0.139]* | 0.072 [0.027 - 0.123]* | 0.062 [0.021 - 0.107]* | 0.070 [0.030 - 0.111]* |
| Scenario 2 ($V_A$=0.3) | 0.063 [0 - 0.215]* | 0.037 [0 - 0.169]* | 0.066 [0.007 - 0.166]* | 0.028 [0 - 0.089]* | 0.052 [0.016 - 0.105]* | 0.043 [0.003 - 0.094]* |
| Scenario 2 ($V_A$=0.1; improper param.) | 0.154 [0.047 - 0.265]* | 0.141 [0.078 - 0.238]* | 0.123 [0.061 - 0.196]* | 0.092 [0.054 - 0.130]* | 0.106 [0.066 - 0.155]* | 0.125 [0.089 - 0.166]* |
| Scenario 2 ($V_A$=0.3; improper param.) | 0.145 [0.025 - 0.269]* | 0.127 [0.044 - 0.222]* | 0.120 [0.049 - 0.202]* | 0.093 [0.042 - 0.143]* | 0.098 [0.045 - 0.157]* | 0.103 [0.059 - 0.151]* |

| | | | | | | |
|---|---|---|---|---|---|---|
| Scenario 3 ($V_A$=0.1; $r_g$=0.1) | 0.013 [0 - 0.088] | 0.009 [0 - 0.045] | 0.013 [0 - 0.064] | 0.004 [0 - 0.042] | 0.004 [0 -0.039] | 0.003 [0 - 0.028] |
| Scenario 3 ($V_A$=0.3; $r_g$=0.1) | 0.015 [0 - 0.151] | 0.019 [0 - 0.091] | 0.007 [0 - 0.078] | 0.011 [0 - 0.059] | 0.007 [0 - 0.049] | 0.008 [0 - 0.046] |
| Scenario 3 ($V_A$=0.1; $r_g$=0.5) | 0.102 [0 - 0.215] | 0.062 [0 - 0.127] | 0.003 [0 - 0.045] | 0.007 [0 - 0.045] | 0.004 [0 - 0.033] | 0.011 [0 - 0.036] |
| Scenario 3 ($V_A$=0.3; $r_g$=0.5) | 0.011 [0 - 0.122] | 0.035 [0 - 0.140] | 0.076 [0 - 0.149] | 0.032 [0 - 0.097] | 0.005 [0 - 0.051] | 0.054 [0 - 0.090] |
| Scenario 3 ($Cov_A$=0.01; $r_g$=0.1) | 0.008 [0 - 0.062] | 0.005 [0 - 0.037] | 0.003 [0 - 0.044] | 0.003 [0 - 0.025] | 0.003 [0 - 0.027] | 0.003 [0 - 0.021] |
| Scenario 3 ($Cov_A$=0.0173; $r_g$=0.1) | 0.072 [0 - 0.133] | 0.005 [0 - 0.069] | 0.005 [0 - 0.052] | 0.024 [0 - 0.046] | 0.004 [0 - 0.030] | 0.005 [0 - 0.036] |
| Scenario 3 ($Cov_A$=0.05; $r_g$=0.5) | 0.009 [0 - 0.074] | 0.016 [0 - 0.067] | 0.014 [0 - 0.043] | 0.005 [0 - 0.027] | 0.009 [0 - 0.034] | 0.005 [0 - 0.029] |
| Scenario 3 ($Cov_A$=0.0866; $r_g$=0.5) | 0.051 [0 - 0.094] | 0.006 [0 - 0.066] | 0.005 [0 - 0.061] | 0.010 [0 - 0.048] | 0.012 [0 - 0.043] | 0.031 [0 - 0.052] |
| Average value (±SD) | 0.0652 (±0.0540) | 0.0625 (±0.0504) | 0.0479 (±0.0466) | 0.0378 (±0.0346) | 0.0358 (±0.0407) | 0.0440 (±0.0435) |

### Figures

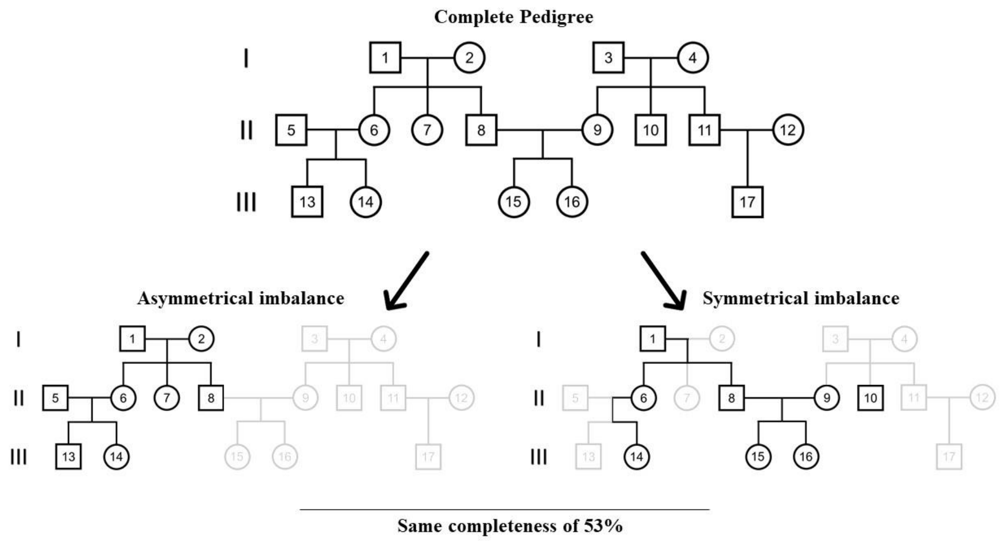


**Figure S1**. Illustration of the two types of pedigrees resulting, for the same degree of completeness, in either an asymmetrical or symmetrical incomplete pedigree.

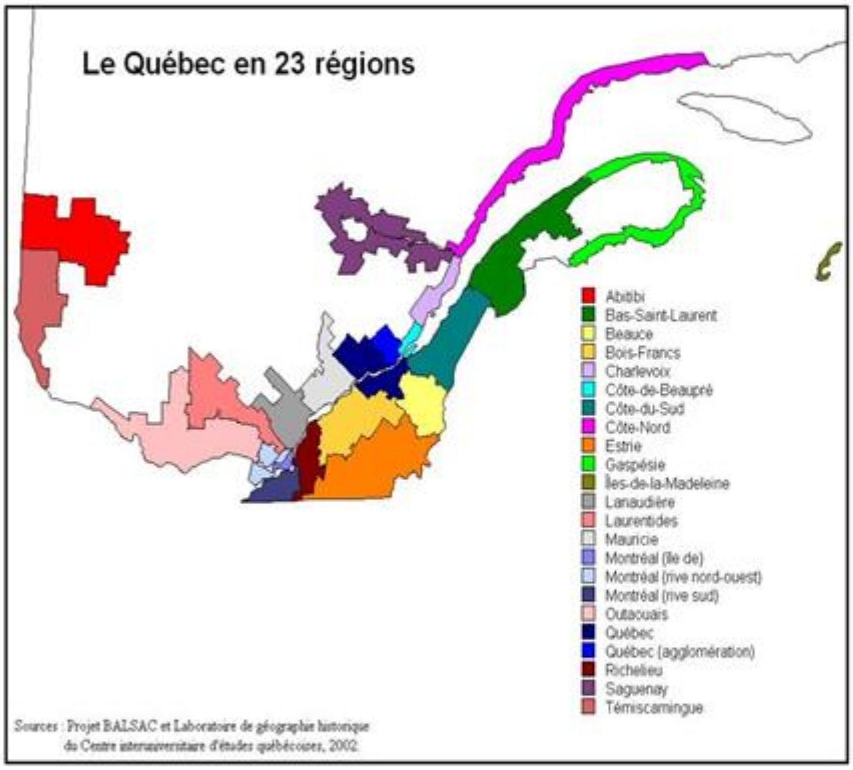


**Figure S2**. Map representing the 23 regions of French-Canada (Québec) for which the BALSAC project contains genealogical records. *Source: Project BALSAC and the laboratory of historical geography at the Inter-University Center for Québec Studies.*

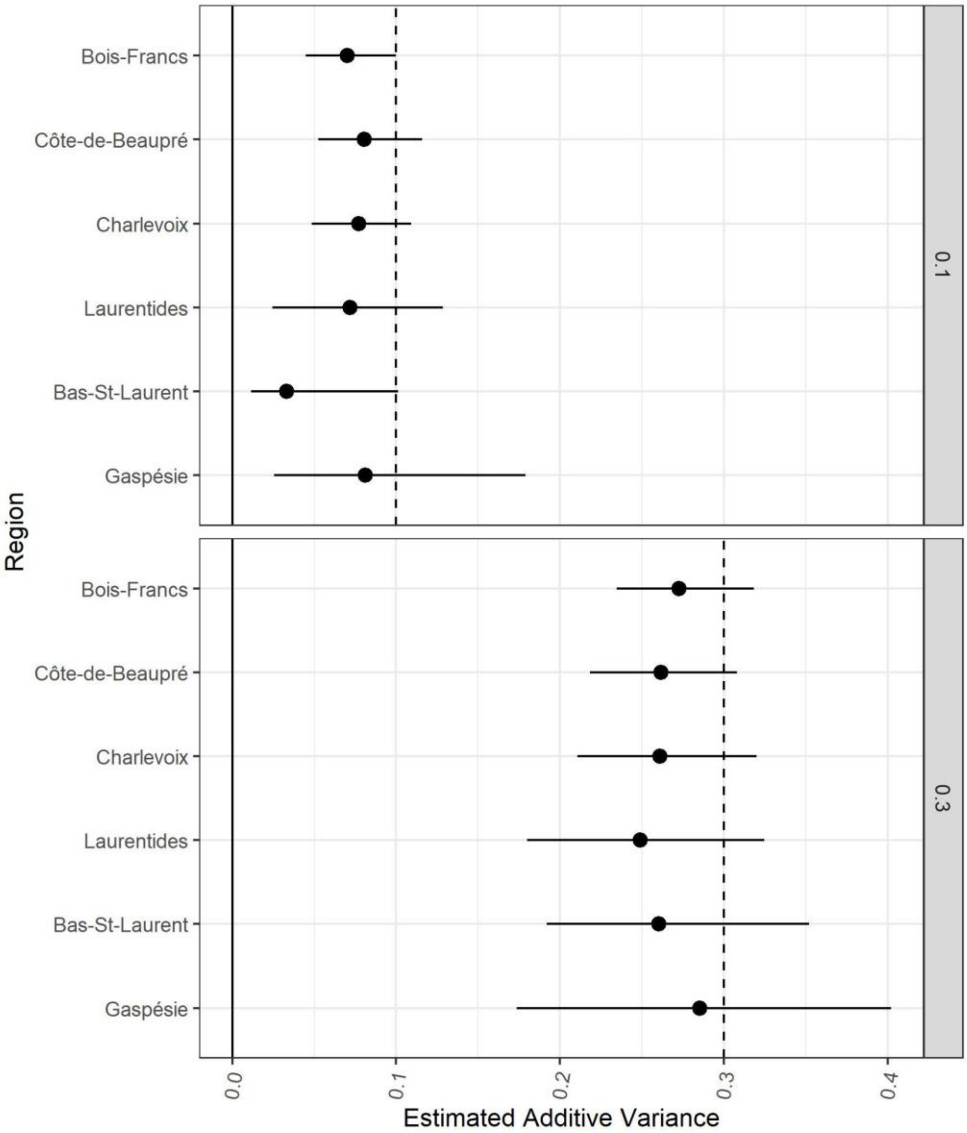


**Figure S3**. Estimates of additive genetic variance for each of the six pedigrees based on Scenario 1 (univariate simple phenotype; $V_A$ = 0.1 or 0.3]) with a genealogical error rate increased to 1%. Each dot represents the average posterior mode-estimates, along with the average 95% credible intervals based on the five simulations per pedigree per parameter value. Vertical dashed lines represent the expected value of the variance.

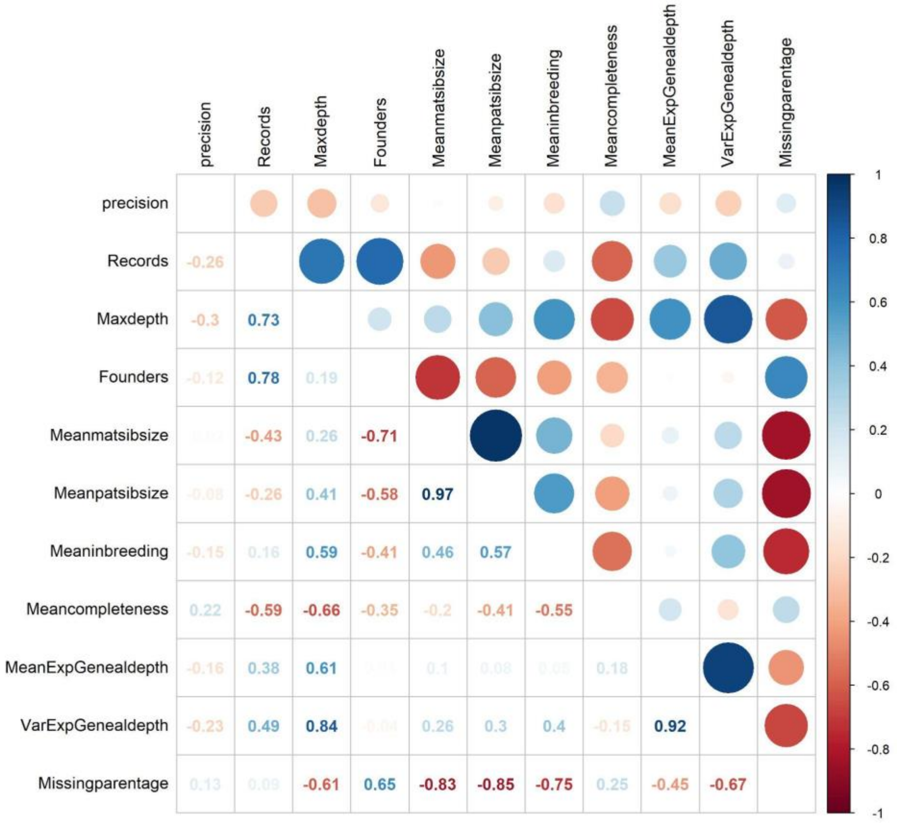


**Figure S4**. Correlation results between different pedigree and genealogical properties and the RMSE-based measure of precision and accuracy.

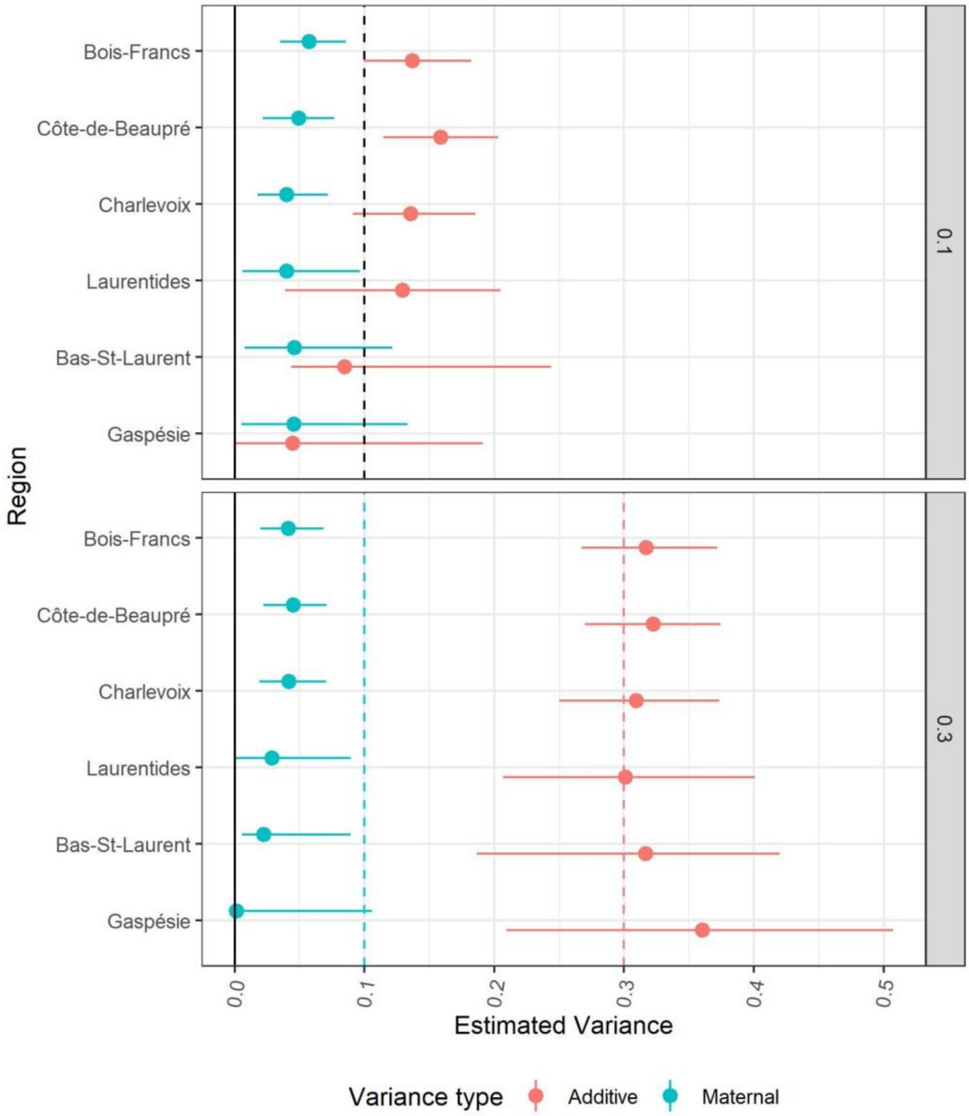


**Figure S5**. Estimates of additive genetic variance and maternal variance for each of the six pedigrees based on Scenario 2 (univariate complex phenotype; $V_A$ = 0.1 or 0,3; $V_M$ = 0.1) with a genealogical error rate increased to 1%. Each dot represents the average posterior mode-estimates, along with the average 95% credible intervals based on the five simulations per pedigree per parameter value. Vertical dashed lines represent the expected value for each variance component.

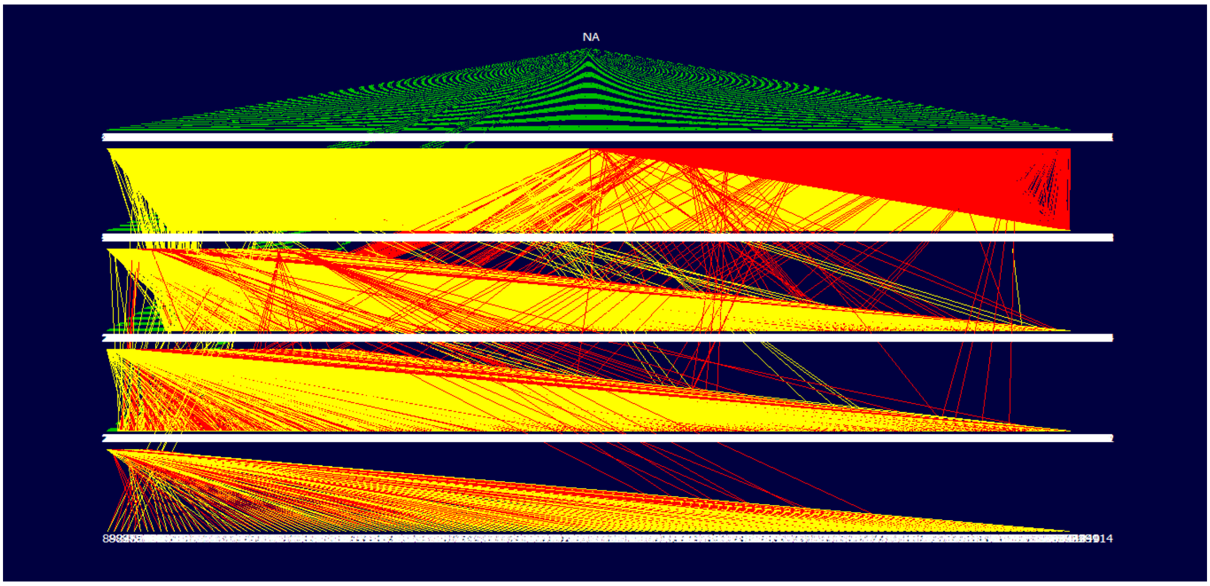


**Figure S6**. Visual representation of the pedigree of the subregion of *Gaspésie*. Red lines correspond to paternal links and yellow lines correspond to maternal links. Green lines correspond to individuals with no known parentage. Drawn using the program *Pedigree Viewer*

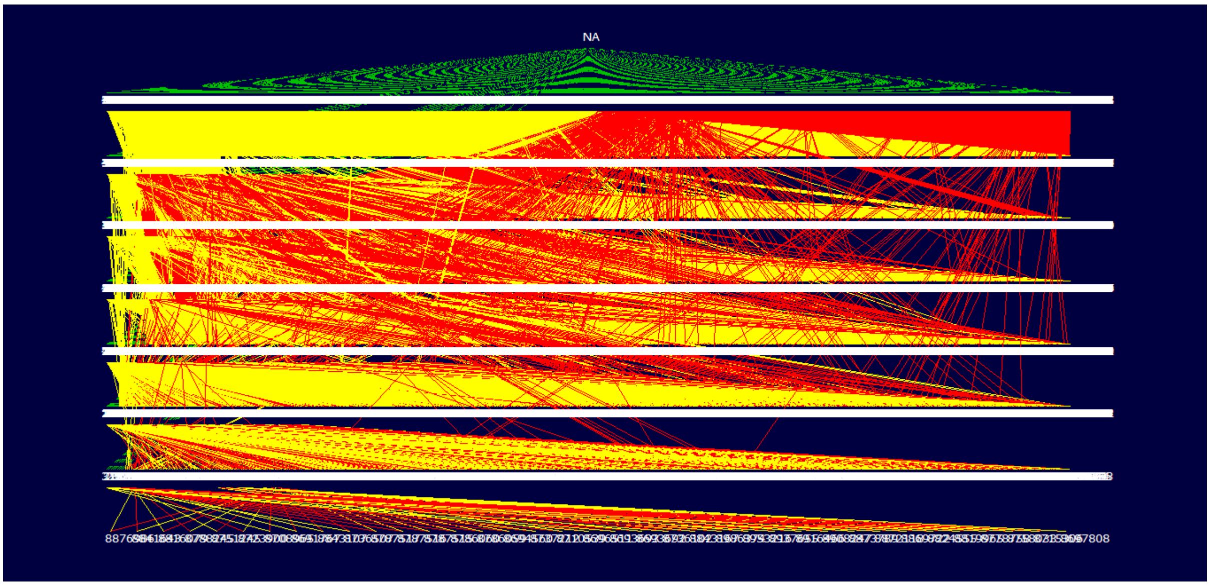


**Figure S7**. Visual representation of the pedigree of the subregion of *Bois-Francs*. Red lines correspond to paternal links and yellow lines correspond to maternal links. Green lines correspond to individuals with no known parentage. Drawn using the program *Pedigree Viewer*
